# A previously reported bottleneck in human ancestry 900 kya is likely a statistical artifact

**DOI:** 10.1101/2024.10.01.615851

**Authors:** Yun Deng, Rasmus Nielsen, Yun S. Song

## Abstract

It was recently reported that a severe ancient bottleneck occurred around 900 thousand years ago in the ancestry of African populations, while this signal is absent in non-African populations. Here, we present evidence to show that this finding is likely a statistical artifact.

---

Recently, Hu et al. [7] identified a severe ancient bottleneck around 900 thousand years ago (kya) in the ancestry of people from Africa. However, they detected no similar evidence of a bottleneck in non-African populations. This is counter-intuitive as the split time between non-Africans and the most closely related African population groups is less than 100 kya [9, 10, 12] and the divergence time between the most diverged African population groups is less than 400 kya [6, 16], and likely considerably less [2, 5, 17]. It should, therefore, not be possible to observe a bottleneck in an African-specific population at 900 kya, and may suggest that some other factor is causing the pattern observed by Hu et al. [7]. In fact, attempts at replicating the bottleneck observation with other methods have failed [19].

The new method, FitCoal, used by Hu et al. [7] to infer the bottleneck is based on fitting the expected Site Frequency Spectrum (SFS) of a demographic model to the observed SFS. The SFS is a very low dimensional compression of the sequence data and SFS-based demography inferences have been known to have identifiability issues [11], namely different population size histories can generate exactly the same expected SFS for an arbitrarily large sample size (number of haplotypes). Even under conditions on the population size function that guarantee identifiability [3], when the observed SFS from a finite number of sites is used in inference, the rate of convergence (as a function of the number of sites) to the true demographic model with a bottleneck is exponentially worse than typical convergence rates for many classical estimation problems in statistics [20]. Furthermore, the space of the expected SFS can be non-convex for a chosen model and if the observed SFS lies outside this set, either due to noise in the observed SFS or model misspecification, then the inferred demographic model can be highly sensitive to small changes in the observed SFS [1, 13]. Given the above facts, it is possible that the inference of a bottleneck made by FitCoal is caused by statistical issues (identifiability, model misspecification, and/or estimation error), and that multiple models, including models that do not include a strong bottleneck, may fit the observed SFS equally well.

To investigate this issue, we first computed the expected SFS under the severe bottleneck model inferred by Hu et al. [7]. We then used *mushi* [4] to infer the best population history whose expected SFS can fit it, with the additional requirement that the population size history does not undergo sudden changes that are too large, thereby disallowing a sharp severe bottleneck as inferred by Hu et al. [7]. More specifically, the trend penalty was set to be (*k, λ*) = (2, 100) and the ridge penalty was set to be 750 in *mushi*. We call this best-fitting model estimated by *mushi* the “synthetic model”. The expected SFS of the synthetic model is very similar to that predicted by the severe bottleneck model of Hu et al. [7] (Figure 1A), showing that models with or without the bottleneck can result in very similar expected SFSs. We then used msprime [8] to simulate 216 haplotypes each of length 100 Mb under the synthetic model (labeled Ground Truth in Figure 1B) to mimic the YRI population data in the 1000 Genomes Project analyzed by Hu et al. [7]. Then, we used Relate [18], MSMC2 [14, 15] and FitCoal [7] to infer population size histories from the simulated data. We note that we provided FitCoal with the exact expected SFS under the synthetic model so that simulation noise has no effect. In the FitCoal analyses, we also eliminated sites with derived allele frequencies between 0.875 and 1, to mimic the truncation procedure used by Hu et al. [7], and used the script provided in their publication to run FitCoal, thereby replicating their inference procedure. All scripts used in these analyses are available at https://github.com/YunDeng98/bottleneck_demography.

**Figure 1.**
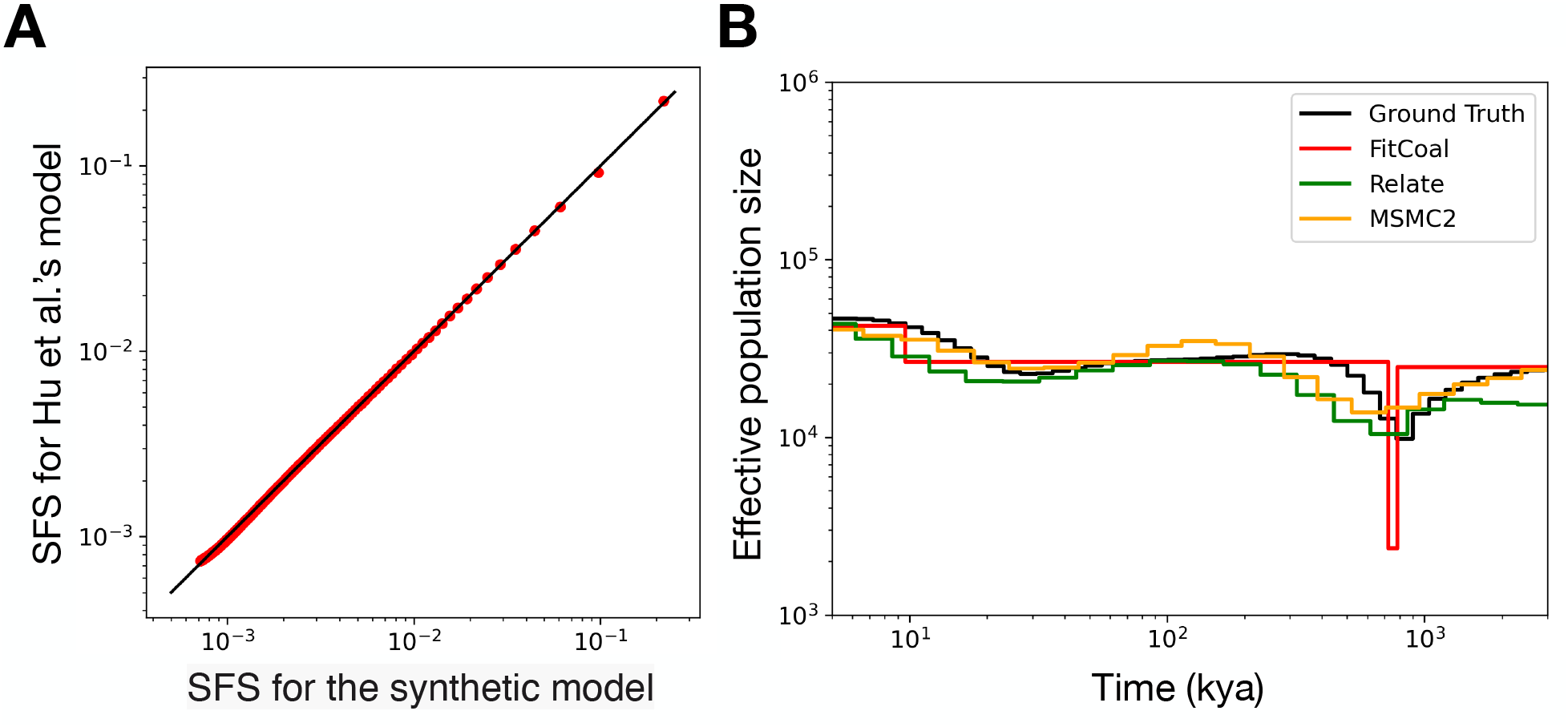
Investigation into the severe bottleneck model. (A) Comparison of the normalized expected SFS under the severe bottleneck model inferred by Hu et al. [7] and that under the synthetic model without an extreme bottleneck. (B) Inferred population size histories for data simulated under the synthetic model (Ground Truth), obtained using Relate, MSMC2 and FitCoal. The exact expected SFS was provided to FitCoal so that simulation noise has no effect, and, as in Hu et al., the generation time was assumed to be 24 years.

The population size histories estimated by the different methods are shown in (Figure 1B). We note that Relate and MSMC2 both estimate population histories roughly similar to that used to simulate the data (i.e., Ground Truth in Figure 1B). However, FitCoal falsely infers a sharp, severe bottleneck instead of the mild population decline assumed in the simulation model. This suggests that under demographic models that generate SFS data similar to that analyzed in Hu et al. [7], FitCoal artifactually tends to infer a sharp bottleneck when there in fact is none. In other words, the reported severe bottleneck is likely a statistical artifact. We note that the original study provided no measures of statistical uncertainty in the inference of the bottleneck. Such measures would likely have shown that there are many demographic models, without a severe bottleneck, that can fit the SFS approximately equally well. Providing valid statistical measures of uncertainty for inferences of demographic models is an important research challenge in computational population genetics.

## Acknowledgments

We thank William DeWitt for helping with using *mushi*. YSS thanks Stephan Schiffels for the helpful discussion on Hu et al.’s work. This research is supported in part by NIH grants R35-GM153400 and R35-GM134922.

